# Transient coupling between subplate and subgranular layers to L1 neurons before and during the critical period

**DOI:** 10.1101/2020.05.05.077784

**Authors:** Xiangying Meng, Yanqing Xu, Joseph P. Y. Kao, Patrick O. Kanold

## Abstract

Cortical layer 1 (L1) contains a diverse population of interneurons which can modulate processing in superficial cortical layers but the intracortical sources of synaptic input to these neurons and how these inputs change over development is unknown. We here investigated the changing intracortical connectivity to L1 in primary auditory cortex (A1) in slices of mouse A1 across development using laser-scanning photostimulation. Before P10 L1 cells receive most excitatory input from within L1, L2/3, L4 and L5/6 as well as the subplate. Excitatory inputs from all layers increase and peak during P10-P16, the peak of the critical period. Inhibitory inputs followed a similar pattern. Functional circuit diversity in L1 emerges after P16. In adult, L1 neurons receive ascending inputs from superficial L2/3 and subgranular L5/6, but only few inputs from L4. A subtype of L1 neurons, NDNF+ neurons, follow a similar pattern, suggesting that transient hyperconnectivity is a universal feature of developing cortical circuits. Our results demonstrate that deep excitatory and superficial inhibitory circuits are tightly linked in early development and might provide a functional scaffold for the layers in between. These results suggest that early thalamic driven spontaneous and sensory activity in subplate can be relayed to L1 from the earliest ages on, that the critical period is characterized by high transient columnar hyperconnectivity, and that in particular circuits originating in L5/6 and subplate might play a key role.

## Introduction

L1 is unique in that most L1 neurons are GABAergic interneurons, showing diversity in firing patterns and expression of molecular markers such as NDNF, 5HT_3_, and others (Hestrin and Armstrong, 1996; Jiang et al., 2013; Lee et al., 2015; Ma et al., 2014; Muralidhar et al., 2013; Schuman et al., 2019; Soda et al., 2003; Tremblay et al., 2016; Winer and Larue, 1989). In adults, L1 neurons can have a profound impact on the sensorily evoked responses of neurons in other layers whose dendritic tufts reside within L1, especially L2/3 cells (Chu et al., 2003; Munoz et al., 2017). L1 is the target of diverse intra-cortical inputs, including crossmodal inputs, as well as subcortical neuromodulatory inputs (Alitto and Dan, 2012; Chu et al., 2003; Ibrahim et al., 2016; Levitt and Moore, 1978; Martin et al., 1989; Mesik et al., 2019; Roth et al., 2016; Zhu and Zhu, 2004). In particular, axonal terminals from subplate neurons are present in L1 (Viswanathan et al., 2017) and stimulation of subplate neurons results in inhibitory responses in Cajal Retzius cells in L1 (Myakhar et al., 2011), suggesting that subplate neurons can directly influence L1 neurons. However, the precise intracortical innervation of L1 neurons and the development of the functional excitatory and inhibitory circuits to L1 neurons is unknown.

The topographic representation of sensory stimulus properties in the cerebral cortex emerges during development, and sensory experience during a critical period can shape this process (Feldman, 2009; Hensch, 2004; Schreiner and Polley, 2014). The underlying circuit mechanisms for the enhanced capacity for plasticity during the critical period have remained unclear. Developing thalamocortical projections first target subplate neurons (Friauf and Shatz, 1991; Hanganu et al., 2002; Higashi et al., 2002; Zhao et al., 2009) and accordingly cortical sensory responses are first observed in subplate neurons (Kanold et al., 2019; Kanold and Luhmann, 2010; Wess et al., 2017). Subplate neurons project to layer 4 (L4) but also to more superficial layers, such as layer 1 (L1) (Deng et al., 2017; Kanold and Luhmann, 2010; Viswanathan et al., 2017; Zhao et al., 2009).

Since during development the pattern of sensorily evoked activity plays a crucial role in shaping cortical organization, L1 neurons are positioned to control activity across the cortical column and might play a key role in processing sensory information and plasticity (Abs et al., 2018). L1 neurons form gap-junction-coupled networks (Chu et al., 2003) and can show synchronous activity in development (Schwartz et al., 1998), suggesting that L1 neurons form co-active networks. Since early network activity can influence cortical development, inputs to L1 have the potential to play a key role in development.

To identify presynaptic inputs to L1 we performed laser-scanning photostimulation (LSPS) combined with whole-cell patch clamp recoding of auditory cortex (A1) L1 neurons in thalamocortical slices from P5 to P32 mice and measured the spatial pattern of excitatory and inhibitory connections. We found that before ear opening (P5-9) L1 neurons receive excitatory input from all layers, especially L2/3 and L5/6. Most L1 neurons (2/3) receive cortical subplate inputs. Inhibitory inputs mostly originated from within L1, L5/6 and subplate. The excitatory inputs from all layers transiently increased at P10-16. In adult, L1 cells receive most excitatory and inhibitory input from within L1, L2/3 as well as superficial L5. Moreover, the functional circuits to L1 neurons diversify during development.

Together, our results indicate that distinct circuit topologies exist during the critical period in rodents. In particular, 1) L1 neurons receive inputs from L4, subgranular L5/6, and subplate, 2) L4 inputs decrease in the second postnatal week, and 3) specific L1 subcircuits emerge in the second postnatal week resulting in circuit sparsification. These findings suggest that subgranular circuits as well as subplate might play a role in controlling activity and plasticity in the superficial cortex before and during the critical period.

## Methods

### Animals

All animal procedures were approved by the University of Maryland Animal Care and Use Committee. Male and female mice (C57Bl/6 background, Jackson Labs) were raised in 12-hr light/ 12-hr dark conditions.

### In vitro Laser-Scanning Photostimulation (LSPS)

LSPS experiments were performed as previously described (Meng et al., 2015; Meng et al., 2019; Meng et al., 2017b; Viswanathan et al., 2017).

### Slice preparation

Mice (C57/Bl6, Jackson Labs; P6-P32) were deeply anesthetized with isofluorane (Halocarbon). A block of brain containing A1 and the medial geniculate nucleus (MGN) was removed and thalamocortical slices (500 μm thick) were cut on a vibrating microtome (Leica) in ice-cold ACSF containing (in mM): 130 NaCl, 3 KCl, 1.25 KH_2_PO_4_, 20 NaHCO_3_, 10 glucose, 1.3 MgSO_4_, 2.5 CaCl_2_ (pH 7.35 – 7.4, in 95%O_2_-5%CO_2_). The cutting angle was ∼15 degrees from the horizontal plane (lateral raised) and A1 was identified as described previously (Cruikshank et al., 2002; Meng et al., 2015; Zhao et al., 2009). Slices were incubated for 1 hr in ACSF at 30°C and then kept at room temperature. Slices were held in a chamber on a fixed-stage microscope (Olympus BX51) for recording and superfused (2-4 ml/min) with high-Mg^2+^ ACSF recording solution at room temperature to reduce spontaneous activity in the slice. The recording solution contained (in mM): 124 NaCl, 5 KCl, 1.23 NaH_2_PO_4_, 26 NaHCO_3_, 10 glucose, 4 MgCl_2_, 4 CaCl_2_. The location of the recording site in A1 was identified by landmarks (Cruikshank et al., 2002; Meng et al., 2014; Meng et al., 2015; Meng et al., 2019; Meng et al., 2017b; Viswanathan et al., 2012; Zhao et al., 2009).

### Electrophysiology

Whole-cell recordings from L1 cells were performed with a patch clamp amplifier (Multiclamp 700B, Axon Instruments) using pipettes with input resistance of 4 – 9 MΩ. Data acquisition was performed with National Instruments AD boards and custom software (Ephus) (Suter et al., 2010), which was written in MATLAB (Mathworks) and adapted to our setup. Voltages were corrected for an estimated junction potential of 10 mV. Electrodes were filled with (in mM): 115 cesium methanesulfonate (CsCH_3_SO_3_), 5 NaF, 10 EGTA, 10 HEPES, 15 CsCl, 3.5 MgATP, 3 QX-314 (pH 7.25, 300 mOsm). Cesium and QX314 block most intrinsic active conductances and thus make the cells electrotonically compact. Biocytin or Neurobiotin (0.5%) was added to the electrode solution as needed. Series resistances were typically 20-25 MΩ. *Photostimulation*: 0.5 – 1 mM caged glutamate (*N*-(6-nitro-7-coumarinylmethyl)-L-glutamate; Ncm-Glu) (Muralidharan et al., 2016) is added to the ACSF. This compound has no effect on neuronal activity without UV light (Muralidharan et al., 2016). UV laser light (500 mW, 355 nm, 1 ms pulses, 100 kHz repetition rate, DPSS) was split by a 33% beam splitter (CVI Melles Griot), attenuated by a Pockels cell (Conoptics), gated with a laser shutter (NM Laser), and coupled into a microscope via scan mirrors (Cambridge Technology) and a dichroic mirror. The laser beam in LSPS enters the slice axially through the objective (Olympus 10×, 0.3NA/water) and has a diameter of < 20 μm. Laser power at the sample is < 25 mW. We typically stimulated up to 30 × 25 sites spaced 40 μm apart, enabling us to probe areas of 1 mm^2^; such dense sampling reduces the influence of potential spontaneous events. Repeated stimulation yielded essentially identical maps. Stimuli were applied at 0.5 – 1 Hz. Analysis was performed essentially as described previously with custom software written in MATLAB (Meng et al., 2014; Meng et al., 2015, 2017a; Meng et al., 2019; Meng et al., 2017b). Activation profiles of neurons across cortical layers during these ages show that LSPS resolution does not vary (Meng et al., 2019). To detect monosynaptically evoked postsynaptic currents (PSCs), we detected PSCs with onsets in an approximately 50-ms window after the stimulation. This window was chosen based on the observed spiking latency under our recording conditions (Meng et al., 2015; Meng et al., 2019; Meng et al., 2017b; Viswanathan et al., 2017). Our recordings are performed at room temperature and in high-Mg^2+^ solution to reduce the probability of polysynaptic inputs. We measured both peak amplitude and transferred charge; transferred charge was measured by integrating the PSC. While the transferred charge might include contributions from multiple events, our prior studies showed a strong correlation between these measures (Meng et al., 2014; Meng et al., 2015; Meng et al., 2019; Viswanathan et al., 2012). Traces containing a short-latency (< 8 ms) ‘direct’ response were discarded from the analysis (black patches in color-coded maps) as were traces that contained longer latency inward currents of long duration (> 50 ms). The short-latency currents could sometimes be seen in locations surrounding (< 100 μm) areas that gave a ‘direct’ response. Occasionally some of the ‘direct’ responses contained evoked synaptic responses that we did not separate out, which leads to an underestimation of local short-range connections. Cells that did not show any large (>100 pA) direct responses were excluded from the analysis as these could be astrocytes. It is likely that the observed PSCs at each stimulus location represent the activity of multiple presynaptic cells. Layer boundaries were determined from the infrared pictures.

Stimulus locations that showed PSC were deemed connected and we derived binary connection maps. We aligned connection maps for each neuron in the population and averaged connection maps to derive a spatial connection probability map. In these maps the value at each stimulus location indicates the fraction of neurons that received input from these stimulus locations. Layer boundaries were determined from the infrared pictures. We derived laminar measures. Input area is calculated as the area within each layer that gave rise to PSCs. Mean charge is the average charge of PSCs from each stimulus location in each layer. Intralaminar integration distance is the extent in the rostro-caudal direction that encompasses connected stimulus locations in each layer. We calculated E/I balance in each layer for measures of input area and strength as (E-I)/(E+I), thus (Area_E_-Area_I_)/(Area_E_+Area_I_), resulting in a number that varied between −1 and 1 with 1 indicating dominant excitation and −1 indicating dominant inhibition.

Spatial connection probability maps show the average connection pattern in each group. To visualize the diversity of connection patterns over the population of neurons in each group we calculated Fano Factor maps. For each responsive spatial location, we calculated the Fano factor of the PSC as σ^2^/μ and plotted maps of the Fano factor.

To compare the large-scale connectivity between cells in each group we calculated the spatial correlation of the binary connection maps in each group by calculating the pairwise cross-correlations (Meng et al., 2019).

### Morphology

Biocytin-filled cells were processed as reported previously (Deng et al., 2017; Meng et al., 2019; Sheikh et al., 2019; Zhao et al., 2009). Cells filled with biocytin were stained and reconstructed (Neurolucida Ver. 2019; MBF Bioscience). Cell morphology was analyzed using the built-in Sholl analysis in Neurolucida.

### Statistics

Results are plotted as means ± s.d. unless otherwise indicated. Populations were compared with a ranksum or Student’s t*-*test (based on Lilliefors test for normality) and deemed significant if p < 0.05.

## Results

### Morphological complexity of L1 neurons decreases over development

We set out to investigate the origin of functional circuits impinging on L1 neurons. Since dendritic complexity is related to the numbers of functional inputs neurons receive, we first investigated the morphological development of L1 cells in A1. We reconstructed 24 L1 cells across different age groups (P10-16: n = 7; P20-23: n = 5; >P28: n = 12) encompassing the critical period for auditory spectral tuning, and through to the mature adult (>P28) (Barkat et al., 2011; Geal-Dor et al., 1993; Willott and Shnerson, 1978; Zhang et al., 2001) (**Fig. 1a**). We then analyzed the dendrites of the reconstructed neurons. This analysis indicated that L1 cells showed the greatest complexity in terms of number of nodes and ends at ∼P10-16 and that this number decreased with age (**Fig. 1b**). Sholl analysis showed that most branch intersections of L1 cells at older age (P20-32) were within ∼100 μm from the cell body while branches were present up to ∼150 μm at P10-16 (**Fig. 1c**). Comparing cells at P20-23 and P28-32 showed that the dendritic branch intersections in the younger group further away from the soma (range from ∼150 to 200 μm) were eliminated. Together, these results indicated that L1 cell morphology became more refined during development.

**Fig. 1.**
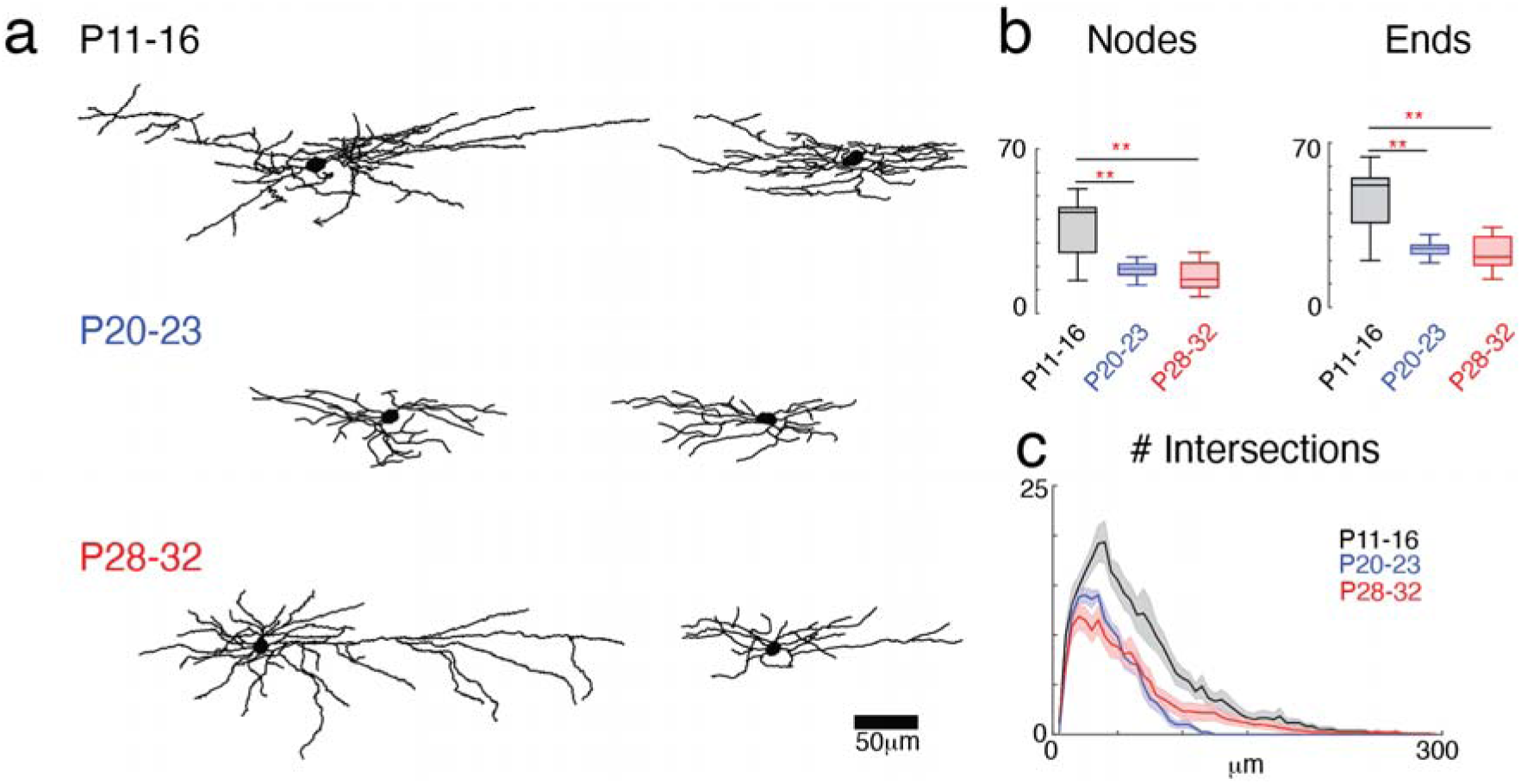
The complexity of L1 cell morphology decreases after P16. (a) Examples of reconstructed L1 cells from different age groups. (b) Total number of nodes and ends of L1 cells at different age groups. Cells in P11-16 group have the most nodes and ends. (c) Sholl analysis of the intersection of sequential serial spheres (5 μm steps) centered on the soma with the cell’s dendrites as a function of the radial distance. The thick line in the middle of the shaded area shows mean across cells and the shaded areas are the standard error.

### L1 neurons receive excitatory inputs from deep layers including subplate and show transient hyperconnectivity of excitatory circuits at P10-16

To investigate the maturation of interlaminar and intralaminar connections to L1 neurons in A1, we applied laser-scanning photostimulation (LSPS) with caged glutamate (Meng et al., 2014; Meng et al., 2015; Meng et al., 2019; Meng et al., 2017b; Shepherd et al., 2003) to map the spatial connectivity of excitatory and inhibitory inputs to L1 neurons in A1 of C57BL/6 mice aged P5-32. LSPS induces action potentials in targeted neurons when the laser beam is close to the soma or proximal dendrites, where a high concentration of glutamate receptors (GluRs) is present. We previously performed cell-attached recordings from cells across the cortical layers at these ages and showed that most photostimulation-evoked APs of L5/6, L4 and L2/3 cells (latencies ≤ 50 ms) were evoked close to the soma at all ages (Meng et al., 2015; Meng et al., 2019; Meng et al., 2017b; Viswanathan et al., 2012), indicating that the spatial resolution of LSPS under our conditions does not vary across age.

To visualize the spatial pattern of excitatory inputs of each cell, we performed whole-cell patch recordings and targeted the laser pulse to multiple distinct stimulus locations. The neurons under the stimulation sites were activated by the laser pulse and generated action potentials. If targeted neurons were connected to the patched cells, we were able to record PSCs. Cells were held at a membrane potential of −70 mV (∼*E*_Cl_) to isolate EPSCs or at 0 mV (∼*E*_glut_) to isolate IPSCs (Meng et al., 2015; Meng et al., 2019; Meng et al., 2017b). The targeted stimulus locations spanned the entire extent of A1, thus enabling us to probe the entire 2-dimensional connectivity pattern of excitatory and inhibitory inputs to a given cell over 1 mm^2^ (**Fig. 2a**). We ensured that the relative laminar locations of sampled cells in each age group were similar (**Fig. 2b**). Activation of glutamate receptors on the cell body and the proximal dendrites caused large-amplitude short-latency direct events (**Fig. 2c**). Since synaptic currents have a distinct latency (> 8 ms), we separated direct and synaptic events using a latency criterion (**Fig. 2c**) (Meng et al., 2015; Meng et al., 2017b).(Meng et al., 2019) The connection strengths were quantified by calculating the PSC charge (Meng et al., 2015; Meng et al., 2019; Meng et al., 2017b) and indicated with pseudocolor scaling in contour maps (**Fig. 2d**). Our recordings showed that L1 cells in A1 received excitatory and inhibitory input from within L2/3 as well as from other layers (**Fig. 2d**).

**Fig. 2.**
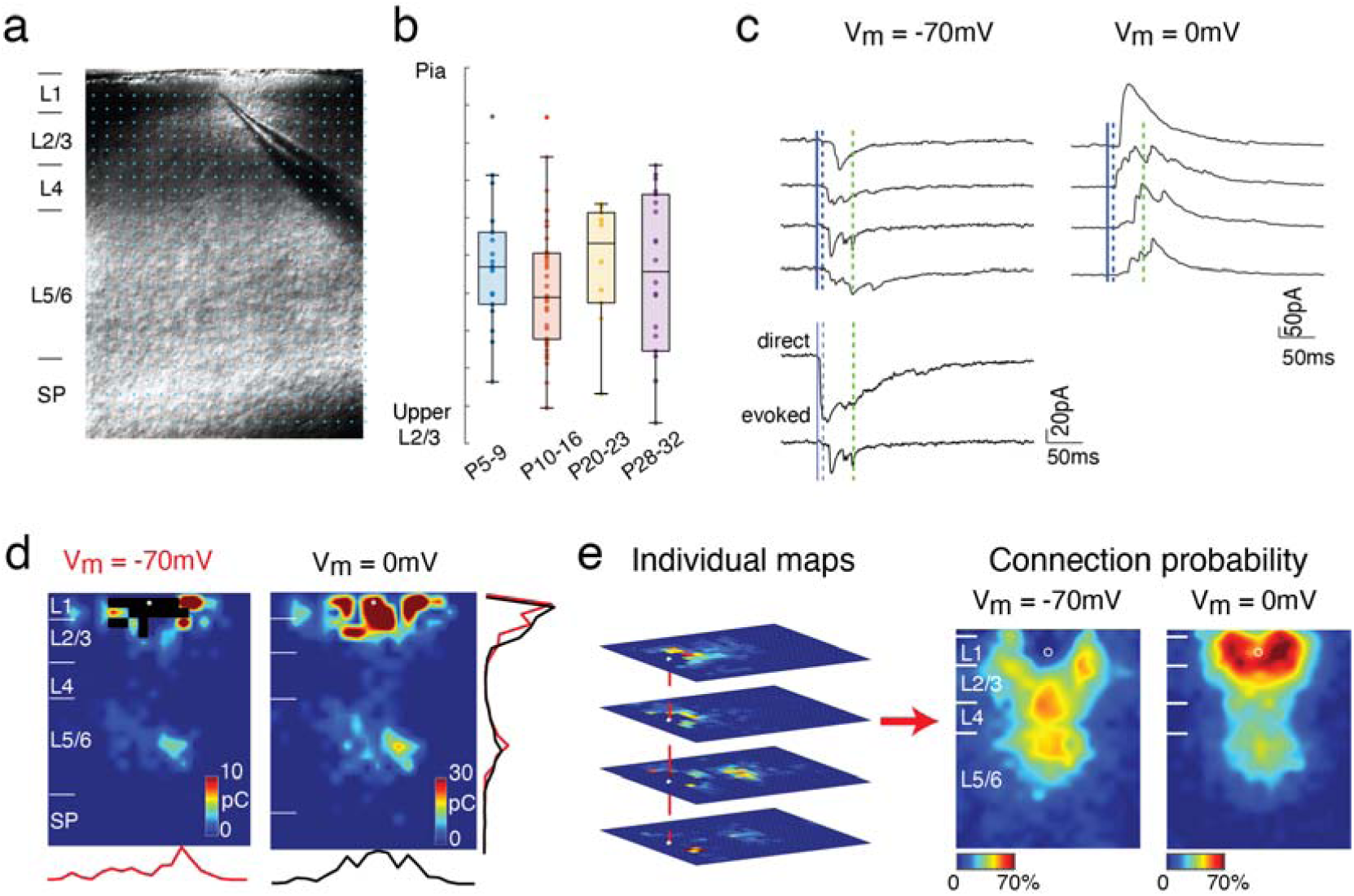
Optical circuit mapping of sources of input to L1 cells. (a) Infrared image of a brain slice with patch pipette on an L1 neuron. Stimulation grids are indicated by the blue dots. The black bars on the left of the image are the layer boundaries (Pia, boundary between L1 and L2/3, boundary between L2/3 and L4, boundary between L4 and L5/6, and the boundary between L6 and subplate). The scale bar represents 100 μm. (b) Relative position of recorded 5cells within L1 for different age groups (P5-9: blue; P10–16: red; P20–23: orange; P28–32: purple). There is no significant difference between different groups (Multicomparison, P5–9 vs. P10–16: P = 0.64; P5–9 vs. P20–23: P = 1; P5–9 vs. P28-32: P = 0.97; P10–16 vs. P20–23: P = 0.77; P10–16 vs. P28–32: P = 0.89; P20–22 vs. P28–32: P = 0.98). (c) Whole-cell voltage clamp recordings with holding potentials at −70 (left) and 0 mV (right) to investigate excitatory and inhibitory synaptic connections, respectively. Shown are example traces evoked by photostimulation at different locations. The solid blue line indicates the time of photostimulation. The dashed blue line marks 8-ms post stimulus, which is the minimal latency for synaptic responses. The dashed green line marks the end of the 50-ms event analysis window. (d) Pseudocolor maps show EPSC (left) and IPSC (right) charge at each stimulus location. White circle indicates the soma location. Horizontal bars indicate layer borders; bar length represents100 μm. (e) illustration of the calculation of the average spatial probability. Input maps are aligned to the soma locations. For each location, the proportion of cells that receive an evoked EPSC or IPSC from that location is calculated and shown with pseudocolor scaling.

We performed LSPS in 90 L1 cells (P6-9: n = 19, P10-P16: n = 38, P20-23: n = 11, P28-32: n =22) from 21 mice (P5-9: n = 6 mice, P10-P16: n = 9 mice, P20-23: n = 2 mice, P28-32: n = 4 mice) and examined the connectivity pattern of excitatory inputs. To visualize connectivity pattern changes over the population of cells, individual LSPS maps were aligned to the cell body position and averaged (Meng et al., 2015; Meng et al., 2019; Meng et al., 2017b); the result is a spatial map of connection probability (**Fig. 2e**). From the averaged maps we observed that L1 cells at all ages received inputs from within L2/3 but also from other cortical layers (**Fig. 2e**). Qualitative analysis of these maps revealed that the spatial profiles changed with development (**Fig. 3a**).

**Fig. 3.**
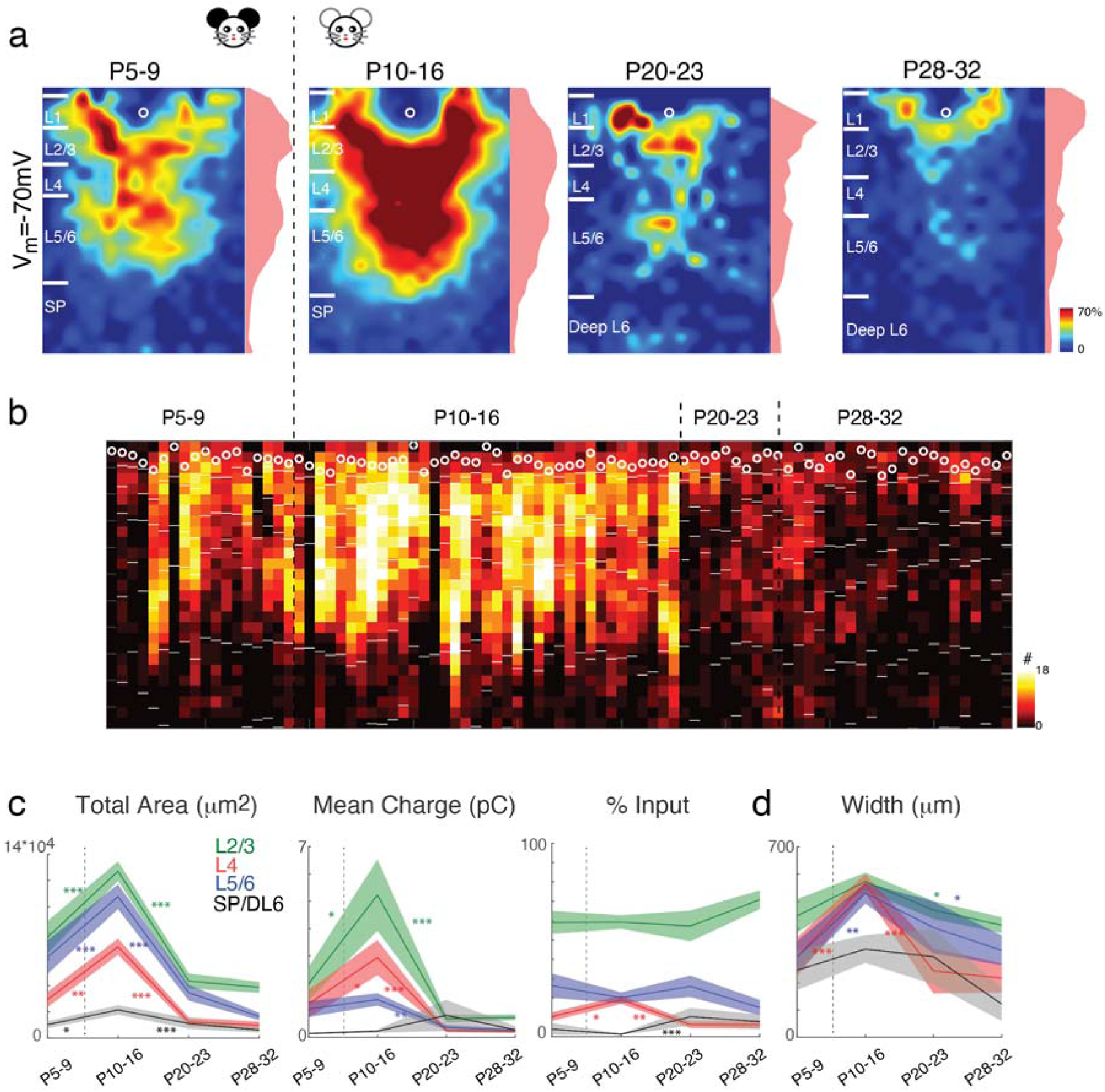
Excitatory circuits to L1 neurons rearrange during development. (a) Average maps of spatial connection probability of excitatory connections in different age groups. The pseudocolor indicates the connection probability. White horizontal bars indicate averaged laminar borders and are 100 μm long. Shaded area on the right side of each probability map indicates the laminar marginal distributions. (b) Summation of input area along laminar direction for all the recorded cells across different age groups. Each column represents one cell. The white circles represent the locations of cell bodies. The short white bars represent layer boundaries. (c) The mean (solid) and SEM (shaded) of total area, mean charge and percentage of inputs L2/3 (including L1, green), L4 (red), L5/6 (blue) and SPN/DL 6 (black) to L1 neurons in different age groups. (d) The mean (solid) and SEM (shaded) of the distance of 80% of excitatory inputs to each L1 cell originating from L2/3 (including L1, green), L4 (red), L5/6 (blue) and SPN/DL 6 (black). The integration distances of L2/3, L4 inputs begin to increase at early age up to P16 and decrease thereafter. P-values are in Supplementary Tables 1-4. The dashed lines in c and d mark the time of ear opening.

To quantify the laminar changes we identified laminar borders for each cell from the DIC images and calculated the input profile from each layer (Meng et al., 2015; Meng et al., 2019; Meng et al., 2017b). To visualize and quantify the differences between cells, we determined for each cell the total area in each layer where stimulation evoked EPSCs in L1 neurons. We find a temporary increase in intracortical synaptic connectivity over development. From the youngest ages on, L1 cells received input from all layers including supragranular, subgranular and subplate layers. Inputs from all layers increased after P5-9, peaked at P10-16 and then decreased (**Fig. 3b, c)**. The laminar distribution of input area for each recorded cell in Fig. 3b shows that the input to L1 cells in adult mice are mainly from intralaminar locations and upper L5. To further quantify the balance of intralaminar to interlaminar inputs, we next determined the relative input L1 neurons receive from each layer. This analysis confirmed that the relative L4 input to L1 neurons reaches its peak at P10-16 (**Fig. 3c**), whereas since the total inputs from L2/3, L4 and L5/6 significantly decreased after P16. In contrast, the relative input from deep L6 increased after P16, peaked at P20, and remained constant until adulthood. Besides the number of inputs the effective synaptic strength contributes to the functional laminar connectivity. To test if synaptic strength from each layer also changed over development, we plotted the mean EPSC charge for EPSCs evoked from each layer (**Fig. 3c**). While the mean synaptic strength for subgranular inputs to L1 remained constant from P5-16, intralaminar and L4 inputs strengthened until P16 (**Fig. 3c**). Input strength decreased after this period in all layers, with the largest decrease occurring in L4 (P10-16 vs. >P28: 525% for L5/6, 1328% for L4 and 711% for L2/3). These results indicate a temporary expansion and strengthening of interlaminar inputs to L1 during the critical period.

So far we assessed circuit expansion across the laminar dimension. Circuit integration across the orthogonal axis in topographic maps indicates topographic integration. Our thalamocortical slices of A1 contain the tonotopic axis, therefore the distance at which presynaptic cells are located in the rostro-caudal axis is a proxy for integration along the frequency axis (Meng et al., 2019; Meng et al., 2017b). We thus compared the rostro-caudal spread or integration distance of inter- and intra-laminar inputs at the different ages (**Fig. 3d**). We find that inputs from L4 and L5/6 originated from a narrow region at P5-9 and that the integration distance increased thereafter, peaking at P10-16. The integration distance in all layers decreased by P28-32 and decreases were largest in L4 (P10-16 vs. >P28 209% for L4 vs. 191% for Deep L6, 150% for L5/6 and 131% for L2/3). Thus, our results indicate that there is a temporary lamina-dependent period of hyperconnectivity of intracortical excitatory circuits in development, which supports integration across the tonotopic axis.

### Transient hyperconnectivity of inhibitory circuits to L1 cells at P10-16

Our data show a substantial remodeling of excitatory connections to L1 cells over development. In L2/3, developmental remodeling of excitatory connections is mirrored by remodeling of inhibitory connections (Meng et al., 2019). We thus investigated if inhibitory connections to L1 also change over development and mapped inhibitory connections by holding cells at 0 mV (**Fig. 2c, d**). Average maps of connection probability across ages appeared different, generally showing a similar developmental trajectory as excitatory connections (**Fig. 4a, b**). This was confirmed quantitatively: the total area generating inhibitory input for all layer decreased after P16 but with modest increase in L2/3 and L4 from P5-9 (**Fig. 4c**, the total inhibitory area of P10-16 vs P5-9 209% for L4, 153% for L2/3, 106% for L5/6, 101% for Deep L6), which is different from excitatory inputs (**Fig. 4c**). The fraction of inhibitory input from L5/6 decreased after P9 and kept constant afterward while the fraction of input from L4 showed modest increase until P16 and then decreased to adulthood. The mean charge of IPSCs originating from within L2/3 and L4 were strongest during P9-P16 and then significantly reduced after that (**Fig. 4c**). In contrast to excitatory inputs, only inhibitory inputs from L4 but not L2/3 and L5/6 showed decreased integration distance with age, indicating a lack of tonotopic refinement (**Fig. 4d**). The marginal distribution of inhibition across tonotopic axis (**Fig. 4a**) eventually become comparable with that of excitation at P28. Thus, together with the changes in excitatory and inhibitory connections, these data show that there is extensive remodeling of excitatory and inhibitory connections to L1 neurons, but that the spatial pattern is distinct for each layer. Our results indicate that there is a temporary increase in relative excitatory but not inhibitory inputs from L4 and L5/6 during the critical period.

**Fig. 4.**
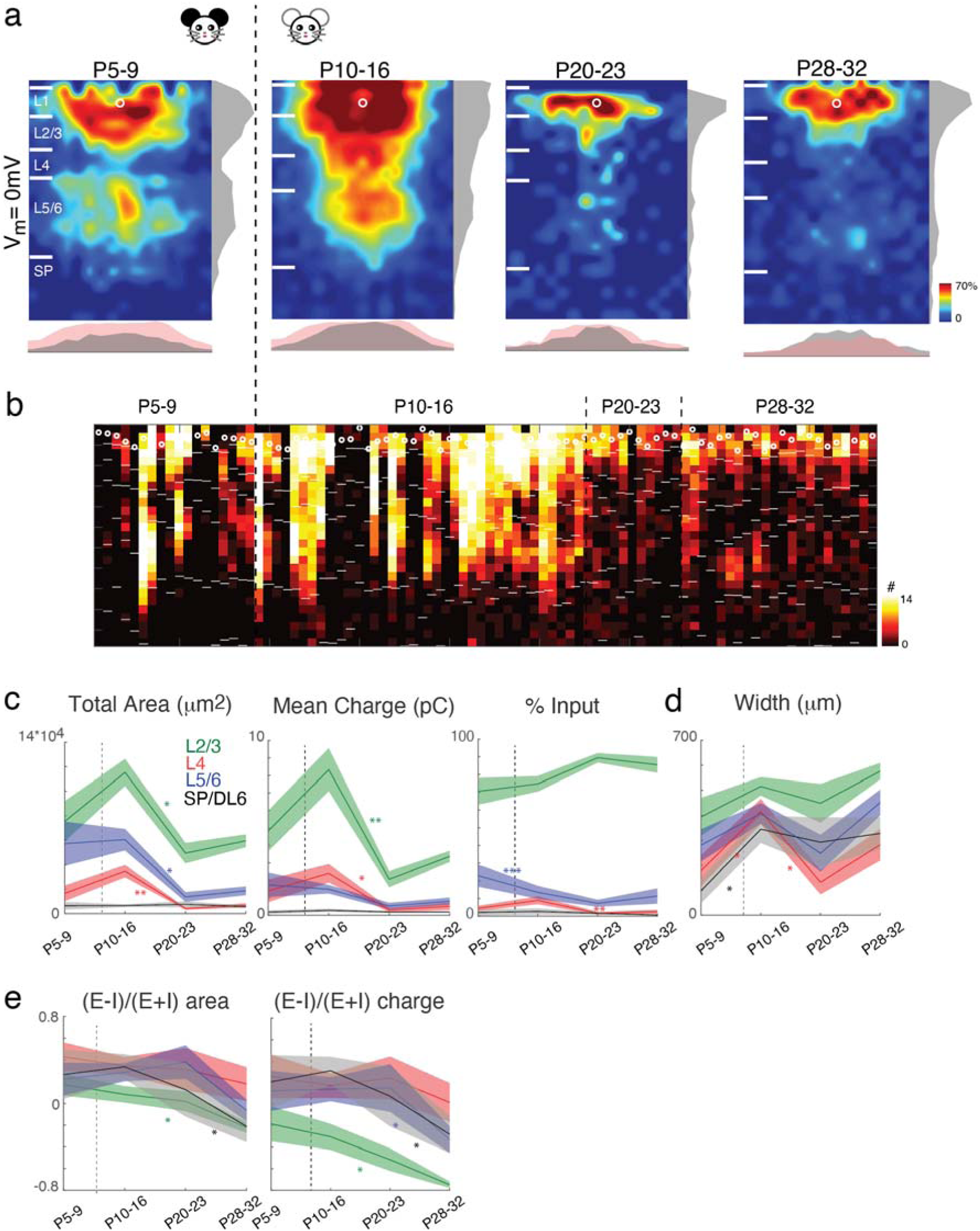
Inhibitory circuits to L1 neurons rearrange during development. (a) Average spatial maps of inhibitory connections to L1 interneurons in different age groups. Pseudocolor represents connection probability. White horizontal bars indicate averaged laminar borders and are 100 μm long. Shaded area on the right side of each probability map indicates the laminar marginal distributions. (b) Summation of input area along laminar direction for all the recorded cells across different age groups. Each column represents one cell. The white circles represent the locations of cell bodies. The short white bars represent layer boundaries. (c) The mean (solid) and SEM (shaded) of total area, mean charge and percentage of inputs from L2/3 (including L1, green), L4 (red), L5/6 (blue) and SPN/DL 6 (black) to L1 neurons in different age groups. P-values are in Supplementary Tables 5-7. (d) The mean (solid) and SEM (shaded) of the distance of 80% of inhibitory inputs to each L1 cell originating from L2/3 (including L1, green), L4 (red), L5/6 (blue) and SPN/DL 6 (black). The integration distances of L4 inputs begin to increase at early age up to P16 and decreases thereafter. P-values are in Supplementary Tables 8. The dashed lines in c and d mark the time of ear opening. (e) Excitatory/inhibitory ((E-I)/(E+I)) balance for inputs from L2/3, L4, and L5/6 based on area or charge. Neurons at adult mice receive more inhibition than excitation, whereas neurons at early age receive more excitation than inhibition. P-values are in Supplementary Tables 9-10.

### E/I balance to L1 cells varies by layer and inhibition in general dominates excitation at P28

Given that both excitatory and inhibitory circuits change over development, and since the spatial and temporal patterns of these changes appeared different, we hypothesized that the balance of excitation and inhibition was changing over development. Studies in the visual cortex have suggested that maturation of functional inhibition, and thus the balance of excitation and inhibition (E/I balance), is a prime driver of critical period plasticity (Hensch, 2004). To determine the functional and spatial balance of excitation and inhibition, we calculated the E/I balance of inputs from each layer based on the area as well as the charge of excitatory and inhibitory inputs (Meng et al., 2015; Meng et al., 2017b) (**Fig. 4e**). This analysis showed that the E/I charge and area ratios are positive for l4 and remained constant until P28 while L2/3, L5/6 and Deep L6 ratios remain constant during P5-P23 and decreased at P28. Thus, E/I balance decreases with age. Inhibition is larger than excitation at adult.

### Emergence of circuit diversity to L1 neurons by P20

L1 neurons form a heterogeneous neuropil in the adult and we thus explored whether this circuit diversity changes over development. Our prior studies of L2/3 circuits showed that circuits to L2/3 could be diverse (Meng et al., 2017b) and that this diversity emerged over development (Meng et al., 2019). We quantified the emergence of L1 circuit diversity by calculating the correlation of the LSPS maps of the sampled L1 cells. We find that correlations for both excitatory and inhibitory maps increased from P5-9 to P10-16, and then decreased by P20-23 and remained constant thereafter (**Fig. 5a**). Thus, L1 neurons form a relatively homogenous population based on intracortical circuit topology during the critical period and circuit diversity emerges after P16.

**Fig. 5.**
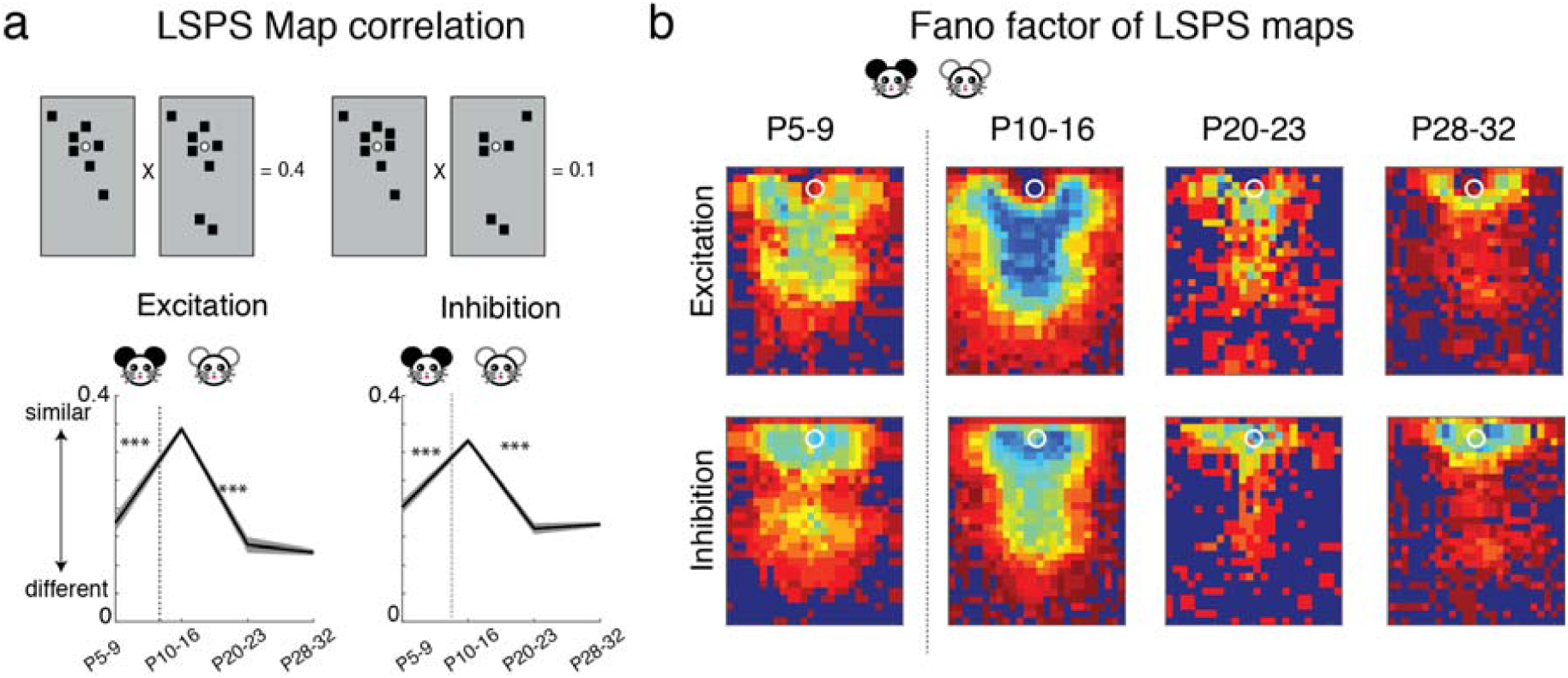
Functional circuit connections to L1 cells become more diverse with age after P16. All maps were aligned based upon cell locations. (a) top: Graphical representation of calculation of pairwise correlation between two maps. Each black square represents the area that has monosynaptic connection to the recorded cell. For the pairwise correlation calculation, we set its value to one and set the area that has no monosynaptic connections to zero. Bottom: the mean (solid) and SEM (shaded) of the pairwise correlations between all excitatory (left) and inhibitory (right) maps. The similarity between connection maps reaches its peak at critical period and decreases after P16. P-values are in Supplementary Table 11. (b) Fano factors (variance/mean) for both excitatory and inhibitory connection maps were calculated based on the response area maps. The Fano factor is represented in pseudocolor. The smaller the Fano factor (blue), the more similarity in the population.

To gain further insight in how inputs from each layer contributed to the circuit diversity we computed the Fano factor for each spatial location over the cell populations in each age group (**Fig. 5b**). We find that the Fano factor was lowest, indicating highest circuit homogeneity, within L2/3 across ages. Interlaminar inputs, especially those from out of column, showed the highest Fano factors, indicating that the increase in circuit diversity is driven by remodeling of interlaminar inputs.

### NDNF+ subtypes of L1 neurons show distinct inputs from subplate at young ages

L1 neurons form a heterogeneous neuropil in the adult and L1 neurons can be grouped by firing patterns and expression of molecular markers such as NDNF, 5HT_3_, and others (Hestrin and Armstrong, 1996; Ma et al., 2014; Muralidhar et al., 2013; Schuman et al., 2019; Soda et al., 2003; Winer and Larue, 1989). We investigated if a molecularly identified subclass of L1 neurons followed a similar trajectory of developmental circuit changes as the general population. We thus crossed NDNF-Cre (Jax #028536) with reporter mice (Jax #007909; CAG promoter-driven TdTomato), patched TdTomato-expressing cells in L1, and performed LSPS (n=70 neurons, P5-9: n =16; P10-P16: n = 18, P20-23: n = 16, P28-32: n = 20) (**Fig. 6a**). At P5-9 NDNF-positive neurons receive strong inputs from L5/6 and subplate rather than the local inputs from within L2/3 (**Fig. 6d**). At P10-16 the interlaminar inputs from L2/3,L5/6 and L4 showed significant increase and then decreased afterwards (**Fig. 6c, d**). At P28, the major excitatory input to NDNF–positive L1 cells are from with L2/3 and L5/6 (**Fig. 6c, d**). Moreover, NDNF-positive L1 cells showed increased circuit heterogeneity after P16 (**Fig. 6e, f**). Inhibitory inputs to NDNF-positive L1 neurons showed a similar pattern. Major inhibitory inputs come from within L2/3 and L5/6 at P5-9. At P10-16 input from L2/3, L4 and L5/6 reached their peaks and then decreased afterward (**Fig. 7a-d**). These results show that NDNF-positive cells receive dominant projections from subplate and L5/6 at the earliest ages and that the circuit changes to NDNF-positive L1 neurons mirror those seen in the general L1 population. Thus, these results suggest that the translaminar hyperconnectivity to L1 neurons at P10-16 is a general property of L1 circuits (**Fig. 7e**).

**Fig. 6.**
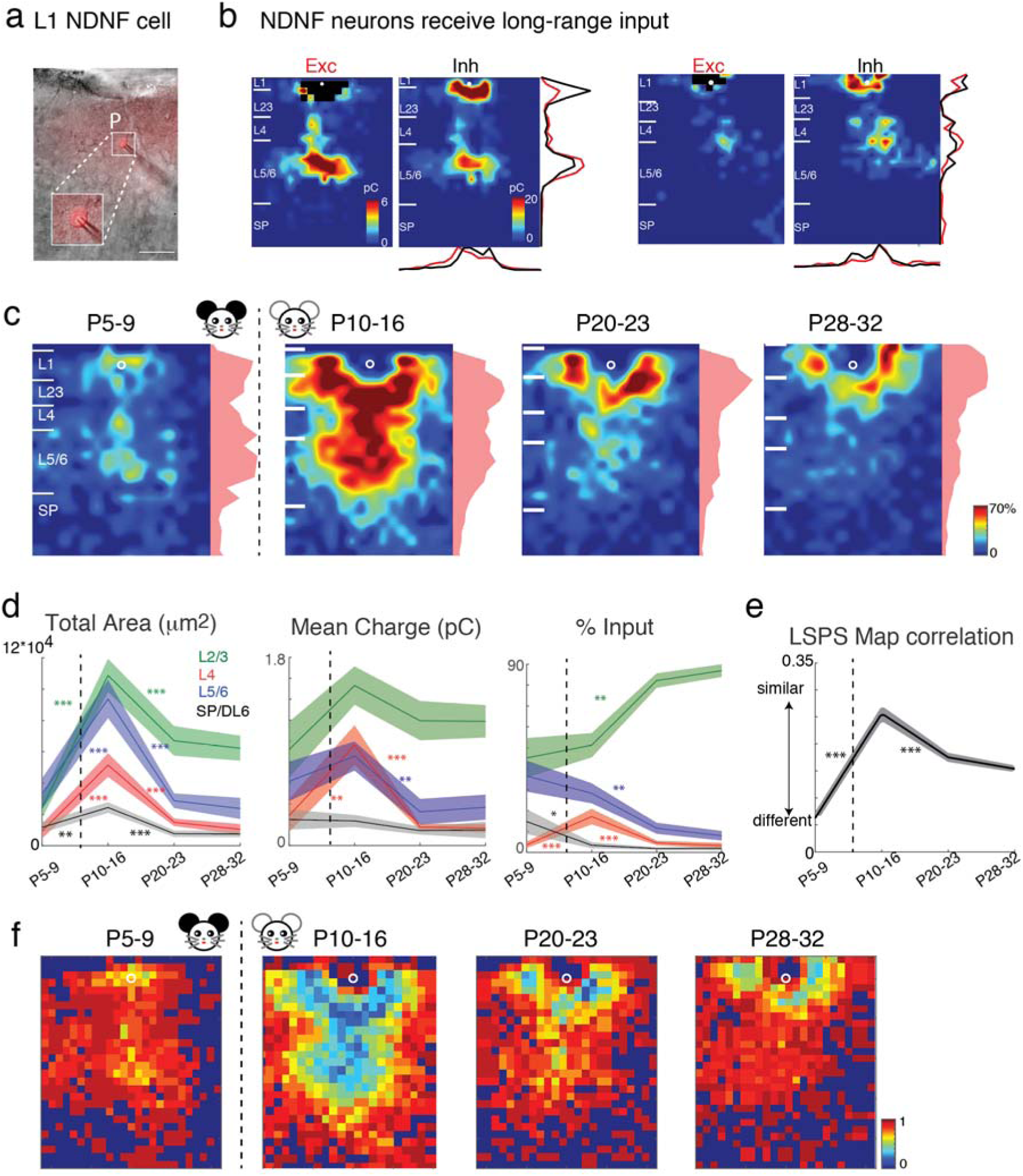
Excitatory connections to L1 NDNF interneurons decrease after P10. (a) In vitro patch clamp recording of NDNF interneuron (red). Imbedded is the twice enlarged area around cell body. Scale bar is 50 μm long. (b) Pseudo color maps show EPSC charge and IPSC charge for two NDNF neurons at P10 (Left) and P12 (Right). (c) Average spatial maps of excitatory connection to L1 NDNF interneurons in different age groups. Pseudocolor represents connection probability. White horizontal bars indicate averaged laminar borders and are 100 μm long. (d) The mean (solid) and SEM (shaded) of total area and percentage of inputs from L2/3 (including L1, green), L4 (red), L5/6 (blue) and SPN/DL 6 (black) to L1 NDNF interneurons in different age groups. At P10-16 L1 NDNF interneurons receive excitatory inputs from all layers. However, in the adult excitatory inputs are mainly from local L1 and L2/3. P-values are in Supplementary Tables 11-14. (e) The mean (solid) and SEM (shaded) of the pairwise correlations between all excitatory maps to L1 NDNF interneurons. (f) Fano factors for excitatory connection maps of NDNF cells were calculated based on the response area maps. P-values are in Supplementary Table18.

**Fig. 7.**
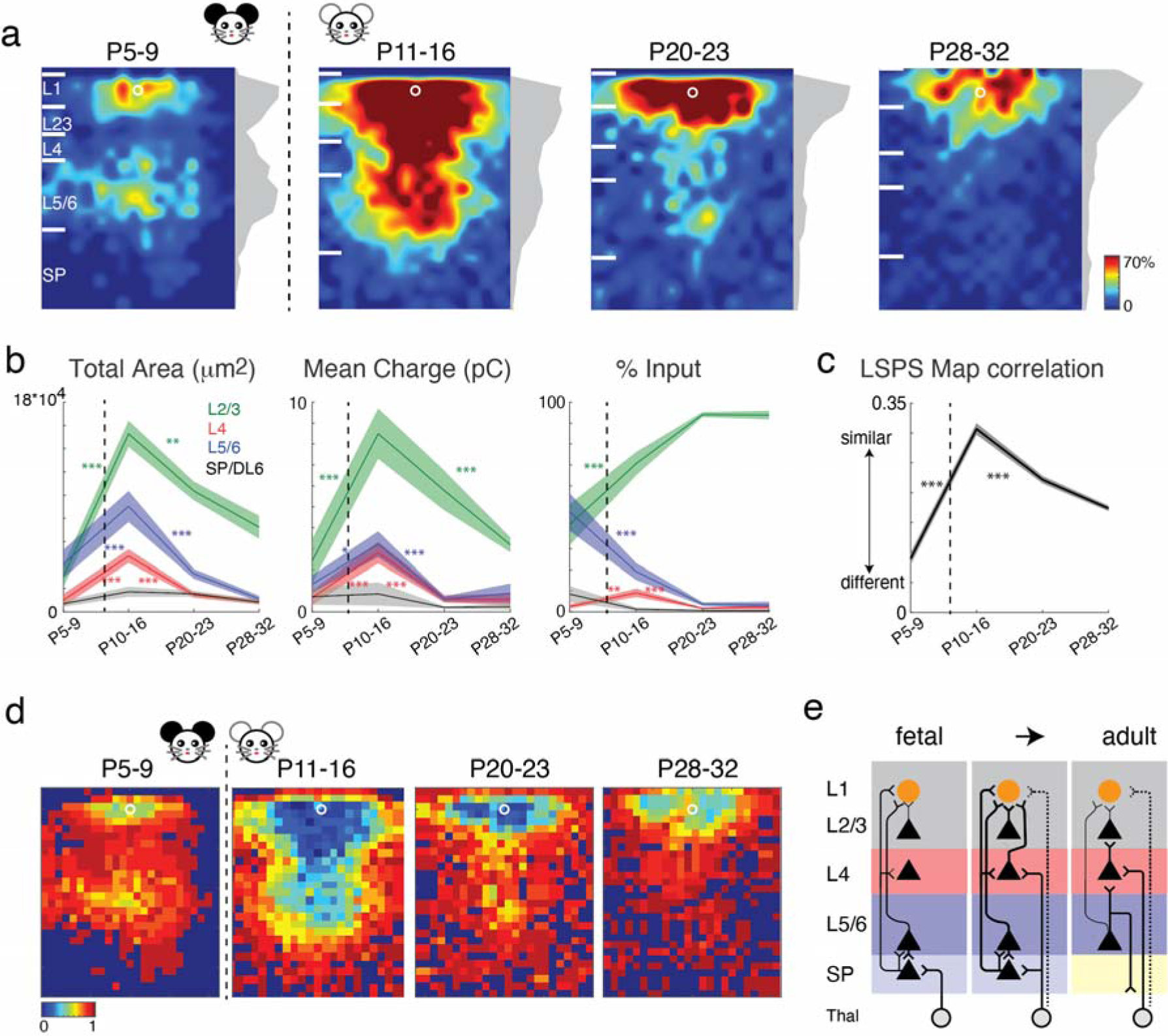
Inhibitory connections to L1 NDNF interneurons decrease after P11. (a) Average spatial maps of inhibitory connections to L1 NDNF interneurons in different age groups. Pseudocolor represents connection probability. White horizontal bars indicate averaged laminar borders and are 100 μm long. (b) The mean (solid) and SEM (shaded) of total area and percentage of inputs from L2/3 (including L1, green), L4 (red), L5/6 (blue) and SPN/DL 6 (black) to L1 NDNF interneurons in different age groups. At P10-16 L1 NDNF interneurons receive excitatory inputs from all layers. However, in the adult inhibitory inputs are mainly from local L1 and L2/3. P-values are in Supplementary Tables 15-17. (c) The mean (solid) and SEM (shaded) of the pairwise correlations between all inhibitory maps to L1 NDNF interneurons. P-values are in Supplementary Table 18 (d) Fano factors for inhibitory connection maps of NDNF cells were calculated based on the response area maps. (e) Cartoon of circuit changes.

## Discussion

In this study we revealed the changing mesoscale inputs to L1 neurons over development. We find that in early development L1 neurons received a large number of excitatory and inhibitory inputs from across all layers of the cortical plate and the subplate. We also find that there is a transient hyperconnectivity of intracortical connections from thalamorecipient layers during the critical period, while in adult animals L1 receives inputs from within L1 and L5. These results suggest that L1 neurons are intimately connected with thalamorecipient layers during the critical period and thus might be able to influence plasticity during the critical period.

We find that dendritic complexity of L1 neurons decreases over development. This is in contrast to the increased dendritic complexity of excitatory neurons in L2/3 (McMullen et al., 1988; Meng et al., 2019), suggesting that dendritic complexity is differentially regulated between neurons.

Thalamic input to the developing cortex first targets neurons deep in the cortex, the subplate neurons (Friauf and Shatz, 1991; Hanganu et al., 2002; Higashi et al., 2005; Kanold and Luhmann, 2010; Zhao et al., 2009) which are the first cortical neurons to respond to sensory stimuli and to show topographic organization (Wess et al., 2017). Subplate neurons send axonal projections to L4 as well as other layers, and excite both excitatory and inhibitory L4 neurons (Deng et al., 2017; Viswanathan et al., 2017; Zhao et al., 2009). Subplate neurons also send axonal projections into the superficial cortex, but the targets of these projections has been unknown. Our results here show that L1 neurons are one target of subplate neurons. While most thalamic projections (e.g. those from MGBv) target L4, corticothalamic projections are also present in L1 (Vasquez-Lopez et al., 2017). Thus, subplate neurons are targeting multiple future thalamorecipient layers.

L1 neurons in adult V1 can respond to visual stimulation as well as locomotion (Mesik et al., 2019), and L1 neurons in S1 can respond to whisker stimulation (Zhu and Zhu, 2004), suggesting that these neurons take part in sensory processing. The short latency of sensory responses in adult L1 neurons suggests direct thalamic activation, but it is unclear when this direct projection emerges. In vitro studies have shown that short-latency responses in cortex first emerge in the subplate before being apparent in L4 (Barkat et al., 2011; Higashi et al., 2002). This, suggests that thalamic activation of L1 neurons also develop after thalamic inputs to subplate have been present. In adult, L1 neurons can relay crossmodal information (Ibrahim et al., 2016; Mesik et al., 2019). Crossmodal information is also present in developing thalamocortical and cortico-cortical projections (Henschke et al., 2018), thus it is possible that L1 neurons in development also receive and relay crossmodal inputs. Since during the critical period L1 neurons receive inputs from the local L4 as well as deep layers, we speculate that L1 neurons initially perform unimodal processing and that the crossmodal processing role of L1 might strengthen after the critical period.

Recent studies have suggested that L1 contains topographic maps (Takesian et al., 2018), which might aid in shaping maps in L4. Since early topographic maps develop in the subplate (Wess et al., 2017), our results here suggest that such maps may result from subplate projections to L1. Studies in cat have implicated subplate neurons for normal critical period plasticity (Kanold and Shatz, 2006). This role of subplate could be due to the existence of subplate inputs across the cortical extent (Viswanathan et al., 2017) coordinating cortical activity and plasticity across layers.

We find that L1 neurons also receive large inputs from L5/6 during the critical period. L5/6 contains multiple cell types with diverse connectivity (Games and Winer, 1988; Kim et al., 2014; Petrof et al., 2012; Prieto and Winer, 1999; Winer and Prieto, 2001). Subclasses of L5/6 cells, e.g. NTSR1-positive L6 neurons, are involved in gain control (Bortone et al., 2014; Guo et al., 2017; Olsen et al., 2012) and we speculate that such a role in development might involve recruitment of L1 circuits which in turn alter the activity of L2/3 pyramidal cells.

L1 neurons comprise a diverse population of neurons (Jiang et al., 2013; Lee et al., 2015; Schuman et al., 2019; Tremblay et al., 2016). Our results show that the overall circuit changes are replicated in at least the NDNF-positive subtypes of L1 neurons which encompasses two classes of L1 neurons (Schuman et al., 2019). Thus, the developmental pattern we delineate here is likely similar for all L1 neurons. While we here utilize NDNF as a marker for a specific class of interneurons, NDNF is a neurotrophic factor that potentially could have effects on target cells (Kuang et al., 2010). While a secretory role for L1 neurons has not been established, a secretory role has been suggested for subplate neurons (Kondo et al., 2015), which express connective-tissue growth factor (CTGF) (Friedrichsen et al., 2003; Heuer et al., 2003; Hoerder-Suabedissen and Molnar, 2013; Kondo et al., 1999; Viswanathan et al., 2012; Viswanathan et al., 2017; Wang et al., 2011). Targets of CTGF and NDNF include neurons and endothelial cells (Khodosevich et al., 2013; Kondo et al., 2015; Ohashi et al., 2014), and we speculate that both deep subplate neurons and superficial L1 neurons form a coupled network in early development that might provide trophic support for neural and non-neural structures in the cortical plate.

We show that L1 neurons receive synaptic inputs from neurons across the cortical column and delineate distinct developmental periods based on the connectivity of L1 cells. In particular, we identify a period of hyperconnectivity from L4 and L5/6 during the critical period, suggesting that these inputs may play an important role before as well during the critical period. We recently showed that there is also a transient period of hyperconnectivity from L5/6 to L2/3 neurons during the critical period (Meng et al., 2019). Therefore, transient hyperconnectivity from L5/6 seem to occur in multiple layers of the cerebral cortex. Thus our data suggests that that transient circuits from subgranular layers L5/6 and subplate might contribute to the plasticity observed during the critical period.

## Acknowledgements

POK, XM designed research. XM performed LSPS experiments and analyzed LSPS data. YX and XM performed morphological analysis. POK supervised research and analyzed data. JPYK contributed reagents. XM and POK wrote the manuscript. All authors edited the manuscript. Supported by NIH R01DC009607 (POK) and NIH R01 GM056481 (JPYK).

