## Supplementary Table for "Transient coupling between subplate and subgranular layers to L1 neurons before and during the critical period"

**Supplementary Tables 1-18**

| Total excitatory area | | | |
| --- | --- | --- | --- |
|  | P5-9 | P10-16 | P20-23 |
| P10-16 | [1.8 × 10^-4^; 1.7 × 10^-5^; 3.9 × 10^-3^; 0.036] |  |  |
| P20-23 | [1.53; 0.41; 0.43; 0.92] | [4.6 × 10^-7^; 1.7 × 10^-5^; 3.9 × 10^-3^; 0.036] |  |
| >P28 | [0.017; 0.049; 0.012; 0.62] | [3.78 × 10^-9^; 3.79 × 10^-9^; 4.26 × 10^-9^; 2.44 × 10^-4^] | [0.99; 0.91; 0.72; 0.34] |

Table 1: P-values for multi-comparison test of total excitatory input area from neurons in L2/3, L4, L5/6 and Deep L6 or Subplate to L1 neurons between different age groups. The p-values for each layer are combined together as vectors: [p-value for L2/3; p-value for L4; p-value for L5/6; p-value for Subplate/Deep L6]

| Mean excitatory charge | | | |
| --- | --- | --- | --- |
|  | P5-9 | P10-16 | P20-23 |
| P10-16 | [0.02; 0.01; 0.24;0.99] |  |  |
| P20-23 | [0.11; 0.68; 0.32;0.07] | [6.3× 10^-5^; 7.0 × 10^-4^; 3.5 × 10^-3^; 0.058] |  |
| >P28 | [0.07; 0.57; 0.15;1.00] | [3.71 × 10^-7^; 1.74 × 10^-5^; 4.20 × 10^-5^; 0.056] | [0.97; 1; 1; 0.056] |

Table 2: P-values for multi-comparison test of mean excitatory input charge from neurons in L2/3, L4, L5/6 and Deep L6 or Subplate to L1 neurons between different age groups. The p-values for each layer are combined together as vectors: [p-value for L2/3; p-value for L4; p-value for L5/6; p-value for Subplate/Deep L6]

| % excitatory charge | | | |
| --- | --- | --- | --- |
|  | P5-9 | P10-16 | P20-23 |
| P10-16 | [1; 0.017; 0.69; 1] |  |  |
| P20-23 | [1; 0.72; 1; 1.46× 10^-4^] | [1; 3.12 × 10^-3^; 0.84; 1.28× 10^-5^] |  |
| >P28 | [0.38; 0.17; 0.28; 0.74] | [0.25; 2.25 × 10^-6^; 0.77; 0.56] | [0.41; 0.91; 0.47; 1.67× 10^-3^] |

Table 3: P-values for multi-comparison test of relative excitatory input from neurons in L2/3, L4, L5/6 and Deep L6 or Subplate to L1 neurons between different age groups. The p-values for each layer are combined together as vectors: [p-value for L2/3; p-value for L4; p-value for L5/6; p-value for Subplate/Deep L6]

| Marginal width of excitatory area | | | |
| --- | --- | --- | --- |
|  | P5-9 | P10-16 | P20-23 |
| P10-16 | [0.11; 9.39 × 10^-5^; 8.0 × 10^-3^;0.81] |  |  |
| P20-23 | [1; 0.94; 0.69; 0.98] | [0.42; 2.4 × 10^-4^; 0.49; 1] |  |
| >P28 | [0.94; 0.72; 0.99; 0.58] | [0.013; 2.87 × 10^-7^; 0.012; 0.074] | [0.87; 1; 0.81;0.45] |

Table 4: P-values for multi-comparison test of marginal width of excitatory input from neurons in L2/3, L4, L5/6 and Deep L6 or Subplate to L1 neurons between different age groups. The p-values for each layer are combined together as vectors: [p-value for L2/3; p-value for L4; p-value for L5/6; p-value for Subplate/Deep L6

| Total inhibitory area | | | |
| --- | --- | --- | --- |
|  | P5-9 | P10-16 | P20-23 |
| P10-16 | [0.091; 0.063; 1; 0.015] |  |  |
| P20-23 | [0.68; 0.59; 0.16; 0.094] | [0.012; 4.9 × 10^-3^; 0.064; 1] |  |
| >P28 | [0.82; 0.51; 0.093; 0.012] | [2.3 × 10^-3^; 1.6 × 10^-4^; 0.014; 0.98] | [0.97; 1; 0.99; 1] |

Table 5: P-values for multi-comparison test of total inhibitory input area from neurons in L2/3, L4, L5/6 and Deep L6 or Subplate to L1 neurons between different age groups. The p-values for each layer are combined together as vectors: [p-value for L2/3; p-value for L4; p-value for L5/6; p-value for Subplate/Deep L6]

| Mean inhibitory charge | | | |
| --- | --- | --- | --- |
|  | P5-9 | P10-16 | P20-23 |
| P10-16 | [0.34; 0.82; 0.51; 0.99] |  |  |
| P20-23 | [0.32; 0.29; 0.13; 0.53] | [6.1× 10^-3^; 0.016; 0.51; 0.47] |  |
| >P28 | [0.41; 0.14; 0.05; 0.53] | [1.2 × 10^-3^; 2.0 × 10^-3^; 0.33; 0.39] | [0.96;0.99; 1; 0.99] |

Table 6: P-values for multi-comparison test of mean inhibitory input charge from neurons in L2/3, L4, L5/6 and Deep L6 or Subplate to L1 neurons between different age groups. The p-values for each layer are combined together as vectors: [p-value for L2/3; p-value for L4; p-value for L5/6; p-value for Subplate/Deep L6]

| % inhibitory charge | | | |
| --- | --- | --- | --- |
|  | P5-9 | P10-16 | P20-23 |
| P10-16 | [0.94; 0.34; 8.6 × 10^-4^; 0.98] |  |  |
| P20-23 | [0.25; 0.63; 0.028; 1.39 × 10^-6^] | [0.37; 0.051; 1; 1.0 × 10^-7^] |  |
| >P28 | [0.13; 0.33; 1.3 × 10^-4^; 0.022] | [0.17; 1.74 × 10^-3^; 0.60; 5.6 × 10^-3^] | [1; 1; 0.73; 2.1 × 10^-3^] |

Table 7: P-values for multi-comparison test of relative inhibitory input from neurons in L2/3, L4, L5/6 and Deep L6 or Subplate to L1 neurons between different age groups. The p-values for each layer are combined together as vectors: [p-value for L2/3; p-value for L4; p-value for L5/6; p-value for Subplate/Deep L6]

| Marginal width of inhibitory area | | | |
| --- | --- | --- | --- |
|  | P5-9 | P10-16 | P20-23 |
| P10-16 | [0.54; 0.043; 0.50; 0.045] |  |  |
| P20-23 | [0.93; 0.97; 0.98; 0.33] | [0.97; 0.044; 0.40; 0.99] |  |
| >P28 | [0.25; 0.81; 0.34; 0.14] | [0.86; 0.23; 0.96; 0.99] | [0.80; 0.62; 0.28; 1] |

Table 8: P-values for multi-comparison test of marginal width of inhibitory input from neurons in L2/3, L4, L5/6 and Deep L6 or Subplate to L1 neurons between different age groups. The p-values for each layer are combined together as vectors: [p-value for L2/3; p-value for L4; p-value for L5/6; p-value for Subplate/Deep L6]

| EI charge: ((E-I)/(E+I)) | | | |
| --- | --- | --- | --- |
|  | P5-9 | P10-16 | P20-23 |
| P10-16 | [0.90; 0.98; 1; 0.98] |  |  |
| P20-23 | [0.51; 1; 1; 0.98] | [0.76; 0.99; 1; 0.86] |  |
| >P28 | [0.021; 0.78; 0.17; 0.32] | [0.031; 0.86; 0.05; 0.04] | [0.74; 0.86; 0.049; 0.045] |

Table 9: P-values for multi-comparison test of EI charge ratio from neurons in L2/3, L4, L5/6 and Deep L6 or Subplate to L1 neurons between different age groups. The p-values for each layer are combined together as vectors: [p-value for L2/3; p-value for L4; p-value for L5/6; p-value for Subplate/Deep L6].

| EI area: ((E-I)/(E+I)) | | | |
| --- | --- | --- | --- |
|  | P5-9 | P10-16 | P20-23 |
| P10-16 | [0.86; 0.97; 0.98; 0.99] |  |  |
| P20-23 | [0.76; 0.96; 0.88; 0.97] | [0.97; 1; 0.96; 0.85] |  |
| >P28 | [0.011; 0.58; 0.31; 0.22] | [0.02; 0.70; 0.054; 0.028] | [0.38; 0.93; 0.12; 0.029] |

Table 10: P-values for multi-comparison test of EI area ratio of L2/3, L4 and L5/6 between different age groups. The p-values for each layer are combined together as vectors: [p-value for L2/3; p-value for L4; p-value for L5/6]

| LSPS Map correlation | | | |
| --- | --- | --- | --- |
|  | P5-9 | P10-16 | P20-23 |
| P10-16 | [3.77× 10^-9^ ; 3.77× 10^-9^ ] |  |  |
| P20-23 | [0.34 ; 0.35] | [3.77× 10^-9^ ; 3.77× 10^-9^ ] |  |
| >P28 | [0.12; 9.3 × 10^-3^] | [3.77× 10^-9^ ; 3.77× 10^-9^ ] | [0.98; 1] |

Table 11: P-values for multi-comparison test of LSPS map correlation between different age groups. The p-values for each layer are combined together as vectors: [p-value for excitatory map correlation; p-value for inhibitory map correlation]

Statistic comparison for NDNF neurons:

| Total excitatory area | | | |
| --- | --- | --- | --- |
|  | P5-9 | P10-16 | P20-23 |
| P10-16 | [1.3 × 10^-8^; 7.0 × 10^-9^; 1.4 × 10^-7^; 6.5 × 10^-3^] |  |  |
| P20-23 | [1.2 × 10^-3^; 0.23; 0.85; 0.46] | [6.5 × 10^-3^; 6.6 × 10^-7^; 6.3 × 10^-7^; 8.2 × 10^-5^] |  |
| >P28 | [2.6 × 10^-3^; 0.97; 1; 0.28] | [1.2 × 10^-3^; 4.93 × 10^-9^; 1.30 × 10^-8^; 5.89 × 10^-6^] | [0.98; 0.37; 0.80; 0.99] |

Table 12: P-values for multi-comparison test of total excitatory input area from neurons in L2/3, L4, L5/6 and Deep L6 or Subplate to NDNF positive L1 neurons between different age groups. The p-values for each layer are combined together as vectors: [p-value for L2/3; p-value for L4; p-value for L5/6; p-value for Subplate/Deep L6]

| Mean excitatory charge | | | |
| --- | --- | --- | --- |
|  | P5-9 | P10-16 | P20-23 |
| P10-16 | [0.50; 2.3 × 10^-3^; 0.96;0.018] |  |  |
| P20-23 | [1; 0.99; 0.035;0.017] | [0.42; 7.46 × 10^-5^; 4.9 × 10^-3^; 1] |  |
| >P28 | [1; 0.99; 0.28; 2.0 × 10^-3^] | [0.38; 3.47 × 10^-5^; 0.073; 0.79] | [1; 1; 0.70; 0.91] |

Table 13: P-values for multi-comparison test of mean excitatory input charge from neurons in L2/3, L4, L5/6 and Deep L6 or Subplate to NDNF positive L1 neurons between different age groups. The p-values for each layer are combined together as vectors: [p-value for L2/3; p-value for L4; p-value for L5/6; p-value for Subplate/Deep L6]

| % excitatory charge | | | |
| --- | --- | --- | --- |
|  | P5-9 | P10-16 | P20-23 |
| P10-16 | [0.89; 1.2 × 10^-4^; 0.60; 0.01] |  |  |
| P20-23 | [2.8 × 10^-4^; 0.90; 2.0 × 10^-3^; 0.013] | [2.0 × 10^-3^; 6.3 × 10^-4^; 0.054; 1] |  |
| >P28 | [9.8 × 10^-5^; 1; 6.6 × 10^-5^; 7.0× 10^-3^] | [9.8 × 10^-5^; 1; 3.1 × 10^-3^; 1] | [0.93; 0.97; 0.76; 1] |

Table 14: P-values for multi-comparison test of relative excitatory input from neurons in L2/3, L4, L5/6 and Deep L6 or Subplate to NDNF positive L1 neurons between different age groups. The p-values for each layer are combined together as vectors: [p-value for L2/3; p-value for L4; p-value for L5/6; p-value for Subplate/Deep L6]

| Total inhibitory area | | | |
| --- | --- | --- | --- |
|  | P5-9 | P10-16 | P20-23 |
| P10-16 | [3.77 × 10^-9^; 3.81 × 10^-9^ 3.68 × 10^-5^; 0.09] |  |  |
| P20-23 | [3.39 × 10^-6^; 0.09; 1; 0.48] | [4.5 × 10^-3^; 1.2 × 10^-7^; 5.42 × 10^-5^; 0.83] |  |
| >P28 | [3.56 × 10^-3^; 0.82; 0.33; 0.99] | [4.3 × 10^-7^; 3.8 × 10^-9^; 1.64 × 10^-6^; 0.14] | [0.11; 0.34; 0.28; 0.62] |

Table 15: P-values for multi-comparison test of total inhibitory input area from neurons in L2/3, L4, L5/6 and Deep L6 or Subplate to NDNF positive L1 neurons between different age groups. The p-values for each layer are combined together as vectors: [p-value for L2/3; p-value for L4; p-value for L5/6; p-value for Subplate/Deep L6]

| Mean inhibitory charge | | | |
| --- | --- | --- | --- |
|  | P5-9 | P10-16 | P20-23 |
| P10-16 | [7.15 × 10^-6^; 3.55 × 10^-4^; 0.034; 0.91] |  |  |
| P20-23 | [6.7× 10^-3^; 1; 0.36; 0.38] | [0.16; 4.17 × 10^-6^; 3.12 × 10^-5^; 0.012] |  |
| >P28 | [0.77; 0.76; 0.23; 0.11] | [1.67 × 10^-5^; 1.12 × 10^-6^; 3.12 × 10^-5^; 0.012] | [0.031;0.63; 0.99; 0.87] |

Table 16: P-values for multi-comparison test of mean inhibitory input charge from neurons in L2/3, L4, L5/6 and Deep L6 or Subplate to NDNF positive L1 neurons between different age groups. The p-values for each layer are combined together as vectors: [p-value for L2/3; p-value for L4; p-value for L5/6; p-value for Subplate/Deep L6]

| % inhibitory charge | | | |
| --- | --- | --- | --- |
|  | P5-9 | P10-16 | P20-23 |
| P10-16 | [4.12 × 10^-4^; 1.9× 10^-3^; 8.4 × 10^-4^; 0.84] |  |  |
| P20-23 | [6.34 × 10^-8^; 0.95; 3.65 × 10^-7^; 0.013] | [0.074; 3.4 × 10^-4^; 0.14; 0.04] |  |
| >P28 | [1.91 × 10^-8^; 0.61; 7.23 × 10^-8^; 0.86] | [0.057; 2.64 × 10^-5^; 0.058; 1] | [1; 0.90; 0.99; 0.03] |

Table 17: P-values for multi-comparison test of relative inhibitory input from neurons in L2/3, L4, L5/6 and Deep L6 or Subplate to NDNF positive L1 neurons between different age groups. The p-values for each layer are combined together as vectors: [p-value for L2/3; p-value for L4; p-value for L5/6; p-value for Subplate/Deep L6]

| LSPS Map correlation | | | |
| --- | --- | --- | --- |
|  | P5-9 | P10-16 | P20-23 |
| P10-16 | [3.77× 10^-9^ ; 3.77× 10^-9^ ] |  |  |
| P20-23 | [3.77× 10^-9^; 3.77× 10^-9^] | [6.59 × 10^-8^ ; 3.77× 10^-9^ ] |  |
| >P28 | [3.77× 10^-9^; 3.77× 10^-9^] | [3.77× 10^-9^ ; 3.77× 10^-9^ ] | [0.37; 8.48× 10^-6^] |

Table 18: P-values for multi-comparison test of LSPS map correlation of NDNF positive L1 neurons between different age groups. The p-values for each layer are combined together as vectors: [p-value for excitatory map correlation; p-value for inhibitory map correlation]
